# Intelligent differential ion mobility spectrometry (iDMS): A deep neural network that predicts optimal space-resolved ion mobility parameters for isomeric monoglycosphingolipids

**DOI:** 10.64898/2026.08.26.747394

**Authors:** Thao Nguyen-Tran, Xun Xun Shi, Emily Hashimoto-Roth, Michael G. Organ, Mathieu Lavallée-Adam, Theodore J. Perkins, Steffany A.L. Bennett

## Abstract

Simultaneous quantification of monoglycosphingolipid stereoisomers is required to monitor changes in defective enzymatic pathways linked to diseases such as Gaucher Disease, Parkinson’s Disease, and Krabbe Disease. Resolution of β-glucosyl and β-galactosyl epimers cannot be achieved by standard liquid chromatography, electrospray ionization, tandem mass spectrometry (LC-ESI-MS/MS). Separation becomes possible when field asymmetric ion mobility spectrometry (FAIMS), also known as differential mobility mass spectrometry (DMS), is added as an orthogonal separation technique to LC. FAIMS/DMS separates epimeric ion clusters in a high versus low electric field (separation voltage, SV) then redirects the target epimeric ions to the mass spectrometer through the application of a direct current (compensation voltage, CoV). Resolving SVs and CoVs must be manually determined for each lipid. Manual derivation is a labour-intensive process that requires pure synthetic standards, limiting the number of stereoisomers a user can include in an assay. To address this problem, we introduce here intelligent DMS (iDMS). iDMS is an *in silico* supervised neural network model that learns the ion mobility relationships between SV and CoV and the monoglycosphingolipid structural features of sugar headgroup, *N*-acyl chain length, and *N*-acyl degree of unsaturation. iDMS predicts the SV and CoV combinations capable of resolving any stereoisomer pair from a training dataset of composed of measured signal intensities across a range of SVs and CoVs of 12 lipids. This machine learning alternative to manual DMS optimization promises to accelerate the deployment of multiple-reaction-monitoring mode (MRM) RPLC-ESI-DMS-MS/MS assays for the routine and rapid quantification of biologically relevant monoglycosphingolipid stereoisomers.

---

Isomeric monoglycosphingolipids are challenging to quantify by mass spectrometry due to their virtually identical structures. Each lipid is defined at the molecular level by its (1) sphingoid base, (2) epimeric sugar unit (glucose or galactose), (3) anomeric α or β glycosidic linkage, and (4) length and degree of unsaturation of the *N*-acyl chain^1^ (Fig. 1). The only structural differences between stereoisomers (species with the same sphingoid base and *N*-acyl chain compositions) are the equatorial or axial stereochemistry of their 4′-hydroxyl group and the α or β glycosidic linkage of their sugar headgroup (Fig 1). These subtle structural differences cannot be resolved by standard reversed or normal phase liquid chromatographic separation (RPLC or NPLC). As a result, quantification using high-performance liquid chromatography, electrospray ionization, tandem mass spectrometry (LC-ESI-MS/MS) commonly reports monoglycosphingolipid stereoisomer abundances as hexosylceramides (HexCers) or hexosyl-sphingoid bases (HexSphs). The hexosyl nomenclature indicates that the sum of both the glucosyl and galactosyl epimers and anomers are reported.

**Figure 1.**
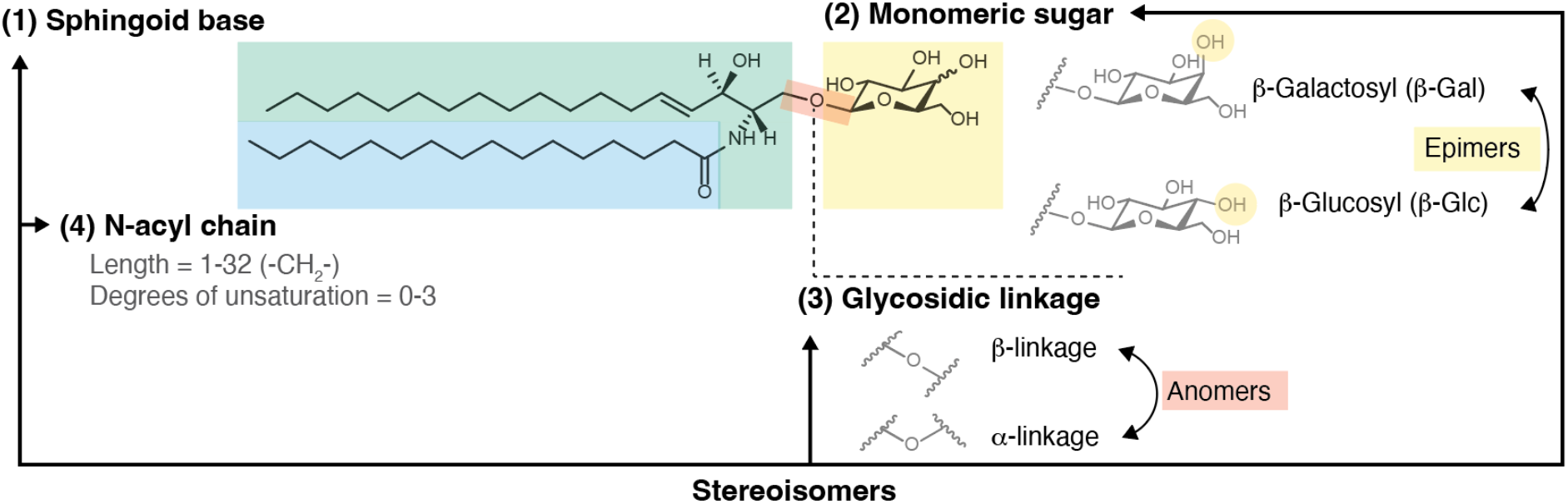
Simple neutral glycosphingolipids. Monoglycosphingolipids are defined by four structural components: (1) their sphingoid base (green), (2) their monomeric sugar unit (yellow), (3) their glycosidic linkage (orange), and (4) their *N*-acyl chain (blue). The sphingoid base can vary in carbon chain length, in the number of hydroxyl groups, and in the number and configuration of double bonds. Sphingoid bases can be further oxidized or methylated. The *N*-acyl chain is a fatty acyl hydrocarbon ranging from one to 32 carbons in length, with varying number of unsaturation, and modifications (hydroxylations, etc.,). Shown are β-GalCer(d18:1/16:0) and β-GlcCer(d18:1/16:0), with (1) a sphingoid base of 18 carbons, two hydroxyl groups (one of which has been replaced by the sugar head-group), and one *trans-*double bond, also known as sphingosine or Sph(d18:1), (2) a galactosyl (Gal) or glucosyl (Glc) monomeric headgroup differing by the stereochemistry of the fourth carbon wherein the hydroxyl group assumes either an axial (Gal) or equatorial (Glc) position (yellow circle), (3) a β-glycosidic linkage differentiating these species from their α anomers, and (4) an *N-*acyl chain of 16 carbons with no degrees of unsaturation (16:0). Stereoisomers refer to monoglycosphingolipids with the same sphingoid base and *N*-acyl chain with either an α or β glycosidic linkage and/or a Gal or Glc headgroup. Epimers are a subset of stereoisomers with the same glycosidic linkage but different sugar headgroups. Anomers are stereoisomers with the same sugar headgroup linked to the sphingoid base by either an α or β glycosidic linkage.

The inability of routine MS/MS approaches to resolve monoglycosylated stereoisomers compromises biological interpretation. Metabolic assessments require quantification of each β-glucosylceramide (β-GlcCers), β-galatosylceramide (β-GalCers), β-glucosylsphingoid bases (β-GlcSphs), and β-galactosylsphingoid bases (β-GalSphs) separately and at the molecular level. For example, homozygous mutations in the *GBA1* gene, encoding the β-glucocerebrosidase enzyme, are the genetic determinants of Gaucher Disease^2^ while heterozygous mutations enhance risk of Parkinson’s Disease^3^. β-glucocerebrosidase hydrolyzes the glucose moiety from β-GlcCers and β-GlcSphs but not the galactose moiety from β-GalCers or β-GalSphs in the absence of cholesterol^4^. Without independent quantification of β-Glc and β-Gal stereoisomers, the effect of mutations on β-glucocerebrosidase enzymatic activity and the *N-*acyl chain substrate specificity cannot be accurately assessed, nor can MS/MS approaches be used to monitor the impact of cause-directed enzyme replacement, substrate reduction, or therapeutic gene replacement. Conversely, mutations in β-galactosylceramidase, encoded by the *GALC* gene, are the genetic determinant of Krabbe disease^5^. β-galactosylceramidase cleaves the galactose moiety from β-GalCers and β-GalSph but not the glucosyl moieties from β-GlcCers or β-GlcSph^6^, further emphasizing the need to efficiently resolve and quantify monoglycosphingolipid stereoisomers for clinical application.

Resolution can be achieved using two advanced separation approaches: hydrophilic interaction chromatography (HILIC)^7^ and differential ion mobility spectrometry (DMS)^8^. HILIC combines the polar stationary phase of NPLC, with a mobile phase resembling RPLC. The polar stationary phase ensures that hydrophilic moieties such as lipids with sugar headgroups are retained on the column until the mobile hydrophobic phase composition is altered. HILIC can efficiently discriminate between β-GlcCer and β-GalCer epimers and β-and α-monoglycosphingolipid anomers but is less effective at quantifying β-GlcCer and β-GalCer stereoisomers^7^. While HILIC ensures that β-GlcCers and β-GalCers elute at different times, all molecular species, defined by their different *N*-acyl chains and sphingoid bases, elute essentially at the same time. This restricted retention range impacts on peak resolution. While it is possible to accommodate a short elution window by reducing the dwell time of each transition to ensure sufficient data points are acquired across all targets for quantification, these shorter dwell times also increase the likelihood of collision-cell cross-talk, reducing accuracy when quantifying lipids in selected or multiple-reaction-monitoring (SRM or MRM) modes. Conversely, if dwell times are not reduced then cycle times are increased, resulting in an insufficient number of data points collected per transition over the short HILIC elution window. Due to these limitations, β-GlcCer and β-GalCer measurements are often reported as combined (total) β- (or α-) GlcCer or β- (or α-) GalCer measurements of all stereoisomers detected using HILIC, biasing quantification towards the most abundant species in each condition and preventing *N-*acyl chain-specific metabolic interrogation.

Alternatively, DMS is a type of Field Asymmetric Ion Mobility Spectrometry (FAIMS) that filters ions according to their different mobilities in a high asymmetric electric field^9^. DMS can resolve neutral glycosphingolipids with different sugar headgroups^8^ by generating ion clusters composed of the same epimers (or anomers) and then separating these clusters based on their interactions with neutral modifier gas molecules (Fig. 2). Because β-GlcCer and β-GalCer stereoisomer ion clusters have different stabilities and different mobilities, each epimeric molecular pair can be resolved by oscillating high and low electric fields within the DMS cell^10^. These fields are generated by an alternating current known as separation voltage (SV). SV is applied perpendicularly to the travelling path of the ions diverting the ions away the mass spectrometer inlet and neutralizing them by collision with the DMS electrodes (Fig. 2). These divergent trajectories can be selectively (and simultaneously) compensated for by application of a direct current or compensation voltage (CoV). CoV opposes SV and re-directs target ion clusters back towards the mass spectrometer intake (Fig. 2). Thus, for any given SV and m/z, a different CoV can be used to resolve each β-Glc and β-Gal stereoisomer at the molecular level. The optimal combinations of SV and CoV allow for the maximal target ion entry at different times. The advantages of DMS to HILIC emerge when DMS is used as an orthogonal separation to RPLC^8^. This combination allows RPLC to separate glycosphingolipids based on their *N*-acyl chain on a timescale of seconds to minutes; DMS to resolve the β-Glc and β-Gal headgroups of each molecular pair within less than a second; and the mass spectrometer in either MRM or SRM mode to quantify individual stereoisomers in milliseconds of scan time (Fig 2). Together, LC-ESI-DMS-MS/MS enables accurate quantification of multiple monoglycosphingolipids in a single method of shorter duration than HILIC with a larger dynamic range^8^.

**Figure 2.**
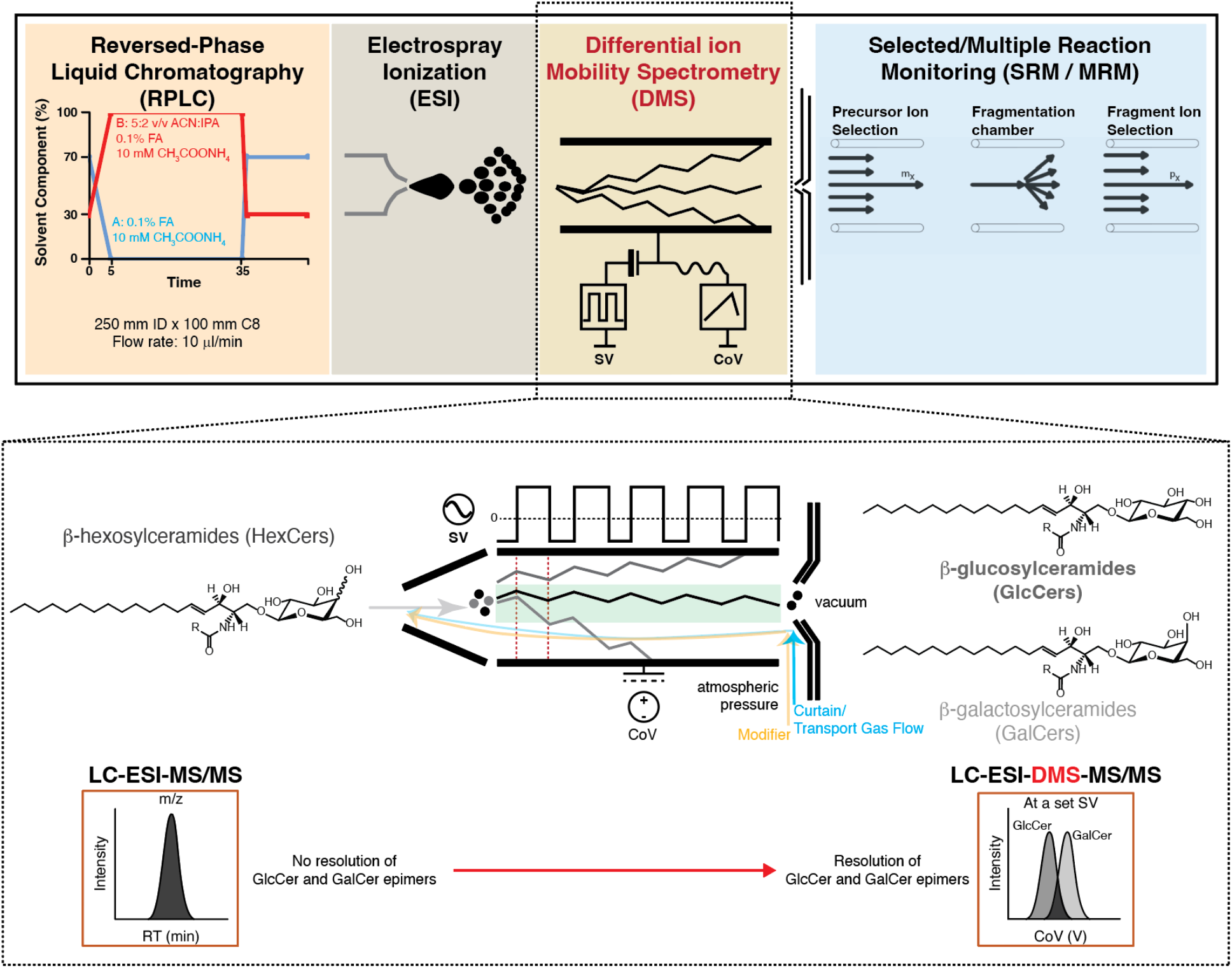
A RPLC-ESI-DMS-MS/MS workflow. (Top panel) Monoglycosphingolipids can be separated at the molecular level by RPLC based on hydrophobicity and ionized by ESI. The sugar epimers can then be orthogonally resolved under atmospheric pressure in the DMS cell, positioned immediately after the ESI source. Stereoisomers are quantified in SRM or MRM mode. **(Inset)** Separation is achieved by the application of an alternating current radio frequency SV that diverts all ions away from the MS intake. This ion-selective diversion is compensated for using direct current CoV, realigning target epimer ions towards the MS intake. Modifier gas is added into the curtain/transport gas flow to increase DMS resolution. Resolution requires unique SVs and CoVs for each Glc and Gal molecular pair. Without DMS, RPLC-ESI-MS/MS can separate molecular isomers with different m/z by their RT but cannot resolve co-eluting epimers with the same m/z (LC-ESI-MS/MS, inset, black peak). With DMS, each epimer eluting at the same RT can be effectively resolved in a single method (LC-ESI-DMS-MS/MS, inset, dark grey and light grey peaks).

The major obstacle to routine LC-ESI-DMS-MS/MS method deployment is the time-consuming, labor-intensive optimization of the compound-dependent FAIMS/DMS parameters. Six DMS machine parameters must be optimized: SV, CoV (at each SV), DMS offset, DMS resolution enhancement, modifier gas composition, and DMS cell temperature. Each parameter influences resolution, peak shape, sensitivity, and reproducibility. The choices of modifier gas, modifier gas composition, DMS resolution enhancement, and DMS cell temperature are epimer- and anomer-dependent but not *N*-acyl chain-specific thus can be determined from a single pair of lipid standards. SV, CoV, and DMS offset however, are compound-dependent, necessitating manual optimization. Manual optimization requires pure standards, limiting the number of lipids a user can optimize to include in a method.

To address these challenges, we present here intelligent Differential Mobility Spectrometry (iDMS). iDMS is a supervised deep neural network with hyperparameters optimized to learn the relationships between three monoglycosphingolipid structural features (sugar headgroup, *N*-acyl chain length, and *N*-acyl degree of unsaturation) and two compound-dependent DMS machine parameters (SV and CoV). iDMS returns to the user the SV and CoV parameter combinations that maximally redirect any target stereoisomer ion into the MS. iDMS is platform-independent. While the neural network was developed using data generated using a SelexION® differential ion mobility device interfaced to a QTRAP 5500 triple quadrupole-linear ion trap mass spectrometer, iDMS accepts training input from any FAIMS-based platform including, but not limited to, Thermo Scientific FAIMS Pro Duo interface^11,12^, custom-built FAIMS instruments such as Heartland Mobility (Bothell, WA, USA)^13^ and Owlstone Inc. FAIMS-chip^11,12,14^. We show here that iDMS can be trained using as few as 6 pairs (12 neutral glycosphingolipid stereoisomers) to achieve separation with a prediction accuracy equivalent to that of empirical derivation. iDMS was developed in Python v3.12.10 using Pandas v2.3.3, NumPy 2.4.0, Tensorflow v2.20.0, and Keras 3.13.0 and is provided as two python scripts and as two Jupyter notebooks freely available at https://github.com/neurolipidomics/iDMS.

## MATERIALS AND METHODS

### Chemicals and Materials

Isopropanol (Optima LC/MS, IPA, A416-4) and LC/MS grade water (Optima LC/MS, W6-4) were purchased from Fisher Scientific Co. Acetonitrile was obtained from J.T. Baker^TM^ (ACN, 9829-03). Ammonium acetate was obtained from Millipore (CH_3_COONH_4_, #2145). Formic acid (FA, LC-MS LiChropur, #5330020050) was purchased from Sigma-Aldrich Canada Co. Ethanol (EtOH) was purchased from Commercial Alcohols (HPLC/LC-MS-high-purity-graded, #P016EAAN). Direct infusion (DI) experiments used a Hamilton Gastight® 1700 series syringe with 500 µl capacity (Hamilton, #81222). T-infusion setup was achieved using three silica capillaries of 120 mm x 75 *μ*m inner diameter (ID) x 363 µm outer diameter (OD) (Polymicro Capillary Tubing, #1068150019), connected by a MicroTee PEEK Assembly for 360 *μ*m OD capillary tubing (Idex Health & Science, #P-888). MicroTight® Adapter PEEK for 1/16” OD tubing to 1/32” OD Tubing (Idex Health & Science, #P881) and red PEEK tubing with 1/16” OD (Sciex, #016316) were used to connect the syringe to the microflow system. A Mini MicroFilter Assembly (Idex Health & Science, M-547) for 1/32” OD with 1 *μ*m porosity and a stainless-steel frit disc was installed immediately before entry to the electrode. Lipid standards were either purchased from, or custom synthesized by, Avanti Polar Lipids as indicated in Supporting Table S1. All lipid parameters standards used in this paper were β-anomers and all had a d18:1 sphingoid base.

### Dataset Acquisition: Direct Infusion-Electrospray Ionization-Differential Mobility Spectrometry-Tandem Mass Spectrometry (DI-ESI-DMS-MS/MS)

Datasets were acquired on a QTRAP 5500 triple quadrupole linear ion trap mass spectrometer (SCIEX) with the SelexION® DMS cell (SCIEX) installed between the orifice and the curtain plate under atmospheric pressure controlled by Analyst 1.6.3 software (SCIEX). Source parameters as well as DMS settings are indicated in Supporting Table S2. Compound-dependent MS parameters were manually optimized in Compound Optimization with DMS mode in Analyst software for each pair of stereoisomers (Supporting Table S3). Previously, we presented a manual DMS optimization workflow for glycosphingolipid stereoisomers and reported IPA as the modifier that achieved optimal separation^8^. Here, we followed this workflow to acquire the training and testing datasets used to develop and validate the iDMS neural network. All lipids were solubilized in EtOH at equimolar concentrations (1 µM) either as a mixture of epimeric pairs or as individual epimers as indicated. Direct infusion (DI) was deployed via T-infusion combining an Agilent Infinity II High-Speed Binary LC Pump (Agilent Technologies, G7120A) and the QTRAP 5500 syringe pump. The syringe pump delivered lipid at a flow rate of 5 µl/min mixed at the T-junction with organic solvent (ACN:IPA at 5:2 v/v, 10 mM CH_3_COONH_4_ and 0.1% FA) delivered by the LC at a flow rate of 10 µl/min. DMS offset (-3.0 V), DMS resolution enhancement (medium, 30 psi), modifier gas composition (low, 173.8 µl/min with IPA as the modifier), and DMS cell temperature (low, 150°C) were as reported previously, optimized using a single pair of epimers (β-Glc(d18:1/16:0)/β-GalCer(d18:1/16:0))^8^. Measured signal intensities were acquired in Compound Optimization mode using Analyst 1.6.3, SCIEX as we have described^8^.

### Preprocessing: Input Dataset Assembly

The iDMS assembly modules “ExtractDMSData_from jdx” and “ExtractDMSData_from mzml” automate the preprocessing of measured signal intensities for parametric modelling and production of iDMS training datasets. Following data acquisition, raw MS files *(.wiff* or other proprietary format) are converted to open-access *.jdx* or *.mzml* format for data parsing. Briefly, for users acquiring training datasets using Analyst 1.6.3, SCIEX, *.wiff* files are converted JCamp files *(.jdx*) using the Analyst 1.6.3 Software Scripts “BatchScript Driver” and “Export to JCamp”^15^. Alternatively, these and other MS proprietary software files can be converted to mzML format (*.mzml*) using Proteowizard^16^. The iDMS preprocessing modules then process and parse the necessary data for parametric modelling from these *.jdx* or *.mzml* files into the format required for iDMS Module 1 to generate the training datasets. The parsing process is accomplished using the iDMS Python scripts or Jupyter notebooks “ExtractDMSData_from jdx” or “ExtractDMSData_from mzml” according to the user’s chosen open-access file format (freely available at https://github.com/neurolipidomics/DMSDataExtractionToolkit). Users of iDMS can follow this processing pipeline (or can use their own data extraction methods to generate their input files by matching the structure of sample *.csv* input files provided (Supporting Files S1, S2). A detailed tutorial of this preprocessing pipeline is provided at (https://www.neurolipidomics.com/DMSDataExtractionToolkit.html).

### Optimization, validation, training, and testing datasets

Datasets were composed of the measured intensities acquired for each monoglycosphingolipid stereoisomer (Supporting Table S1) at 42 SV voltages ranging from 0-4100 in 100 V steps, ramping through 101 CoV voltages from -10 V to 10 V in 0.2 V increments at each SV. Data were acquired as technical replicates of 34 glycosphingolipid stereoisomers, comprising 2856 datasets composed of 101 observations per dataset per lipid. A total of 2352 datasets were used for optimization, validation, and training representing the technical replicates of 28 lipids (14 pairs of lipid epimers infused individually, β-Glc and β-Gal(d18:1 and d18:1/8:0; 14:0; 16:0; 16;1; 18:1; 20:0; 21:0; 22:0; 23:0; 24:0; 24:1; 25:0; 31:0). Testing datasets were composed of 504 datasets from the technical replicates of six lipids infused individually (β-Glc and β-Gal(d18:1/18:0; 22:1; 26:0), held-out from the optimization, validation, and training datasets. Prediction accuracy and resolution were further verified by deploying the iDMS predicted parameters in a RPLC-ESI-DMS-MS/MS MRM workflow quantifying 6 concentrations (0.004-1µM) of custom-synthesized lipids loaded both as mixtures of epimers and individually.

### Module 1: Parametric Modelling

In a typical DMS workflow, the optimal SV and CoV combinations are determined empirically by comparing ionograms where, at each SV, measured signal intensities are expressed as a function of CoV (Fig 3a, Measured). iDMS Module1_Parametric Modelling automates this process, generating both ionograms and training datasets at every SV from the user’s assembled input dataset of measured signal intensities (Fig 3a, Measured, Supporting Table S4). The CoV producing the maximal signal intensity at each SV is extracted (Supporting Table S4). Measured signal intensities are normalized to the maximal signal intensities at each SV (Fig 3a, Normalized, Supporting Table S4). Each normalized ionogram is modelled as a Gaussian curve to establish the µ and σ used for both iDMS training and subsequent assessment of stereoisomer separability (Fig 3a, Gaussian Mode, Supporting Table S4).

**Figure 3.**
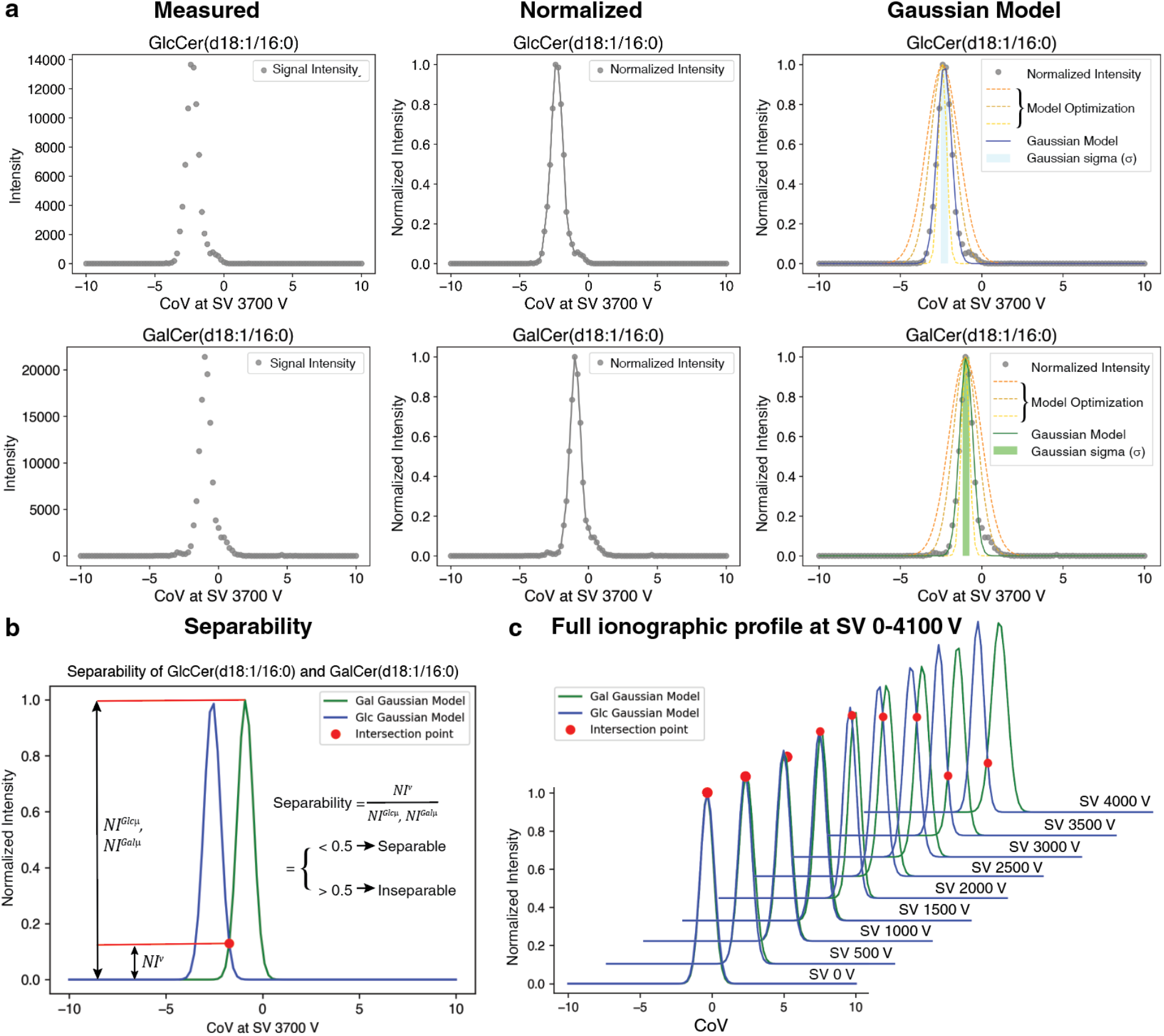
iDMS Module 1: Parametric modelling. **a)** Ionograms plot measured signal intensity, for a given SV, as a function of CoV. iDMS automates ionogram production for all measured signal intensities in the training dataset (Measured). Intensities are normalized to the maximal intensity at each SV (Normalized). The normalized distributions are fitted to their Gaussian function deriving CoV *μ* and σ (at each SV for all monoglycosphingolipid epimers (Gaussian Model). **b)** The Gaussian fits of the Glc and Gal intensity distributions at every SV are plotted to determine their intersection points and separability, defined as the SV and CoV combinations at which the valley intensity (NI^v^) at the intersection point between the two intensity distributions is <0.5 of the normalized signal intensity. **c)** Ionograms produced at each step are returned to the user.

Because µ and σ are derived from normalized signal intensities, the Gaussian function is simply:

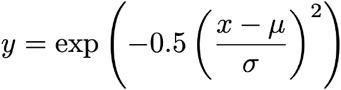

where x= the array of CoV values at a given SV and y=the corresponding array of normalized intensities at each CoV. The Gaussian model fit at every SV and CoV is optimized using Model class in the lmfit package v1.3.4^17^, assigning the initial σ value to 1 and solving for the best fit parameters (Fig. 3a, Gaussian Model, Supporting Table S4). Module 1 assesses overlap of the Glc and Gal stereoisomer ionograms at each SV by solving for the intersection points of both Gaussian distributions and classifying the normalized distributions as separable or inseparable. Separability (resolution) is defined as the SV and CoV combinations at which the valley intensity (NI^v^) at the intersection point between the two intensity distributions is less than 50% of the normalized signal intensity at the Glc µ and Gal µ (NI^Glcµ^, NI^Galµ^) CoV (Fig 3b, Separability, Supporting Table S4).

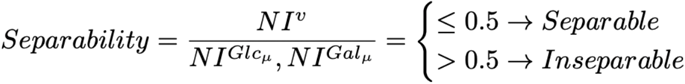

where NI^v^ is the normalized valley intensity and NI^Glcµ^, NI^Galµ^ are the maximal normalized peak intensities at each SV for each stereoisomer.

Module 1 will execute only if the following criteria are met: 1) Training datasets are composed of matched stereoisomer molecular pairs (i.e., pairs of Glc and Gal monoglycosphiongolipids with the same sphingoid base and *N*-acyl molecular identities); 2) The signal intensities are measured across the same CoV range and using the same CoV ramp step (e.g., all stereoisomers in a training set are measured across a CoV range of -10 to 10 V using a 0.2 V step). If one SV is missing, iDMS will perform a quality check on the input training dataset and only keep the SV at which there are DMS measurements for both isomers.

### Module 2: Supervised Learning Problem Formulation of the iDMS Neural Network

Our supervised learning problem was to predict the Gaussian CoV µ (CoV at maximal signal intensity) required to redirect any β-Gal and β-Glc monoglycosphingolipid stereoisomer into the mass spectrometer at all of the SVs provided in the training dataset as a function of three features: 1) *N*-acyl chain length (integer feature), 2) degree of unsaturation of the *N*-acyl chain (integer feature) and 3) SV (integer feature) (Fig. 4a,b) As a result, Module 2 will execute only if the training dataset generated by Module 1 includes a) stereoisomer epimers with at least 2 different *N*-acyl chain lengths and 2 different degrees of unsaturation in the *N-*acyl chain and b) 2 different SVs. Thus, the smallest dataset possible to operate iDMS Module 2 is composed of measured intensities from 4 monoglycosphingolipids (2 pairs of stereoisomers) measured at a minimum of 2 SVs.

**Figure 4.**
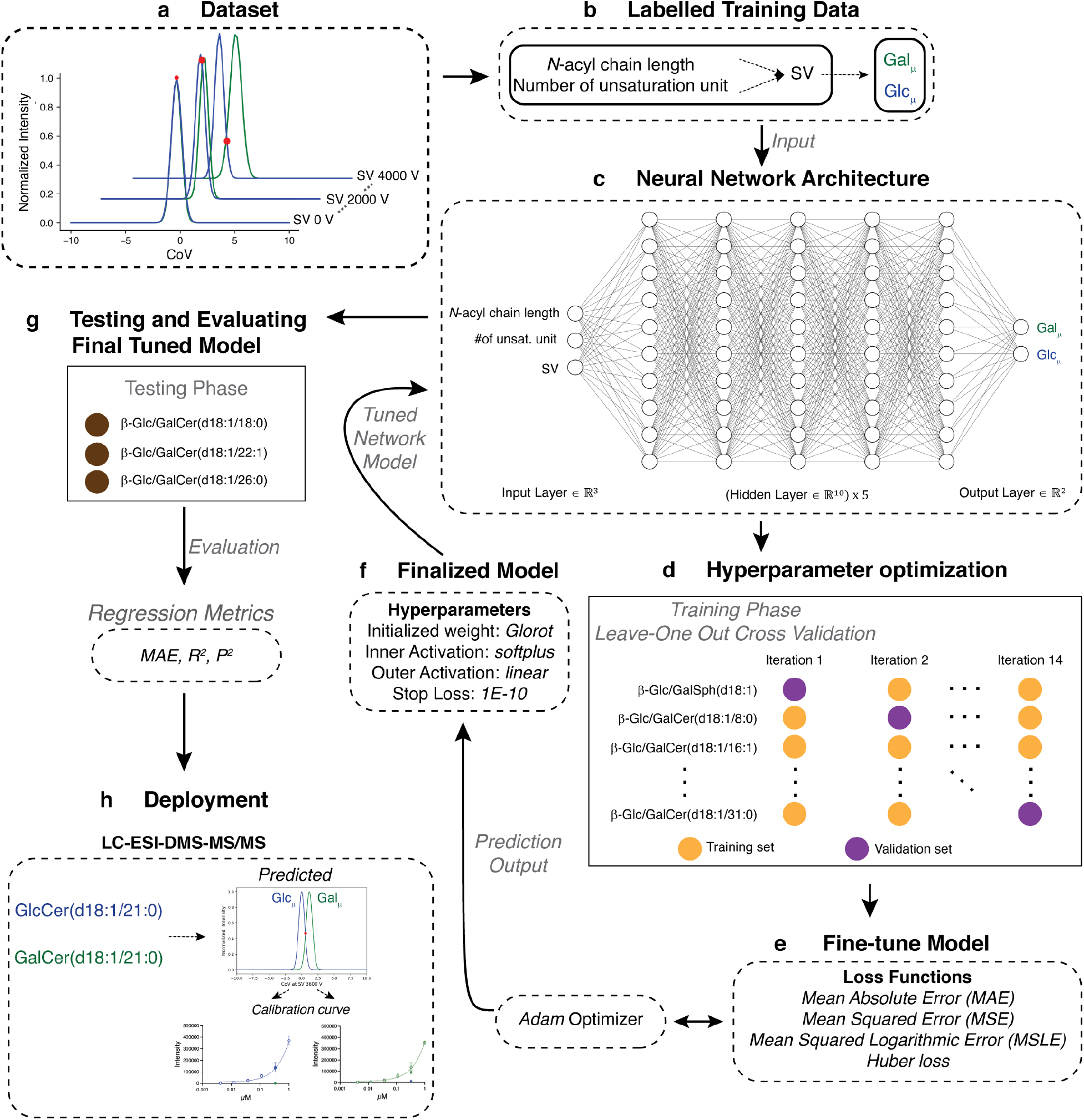
iDMS Neural Network Development and Evaluation. **a)** and **b)** iDMS obtains the Gaussian features (*μ* and σ) and the lipid features (*N-*acyl chain length and the number of unsaturation unit) of each monoglycosphingolipid epimer at corresponding SV from an empirical (training) dataset. **c)** The iDMS neural network is composed of seven interconnected layers (input layer, five hidden layers and output layer). **d)** A leave-one-lipid-pair out cross-validation (LOOV) approach was used to evaluate combinations of hyperparameters during tuning process, according to **e)** mean absolute error (MAE), mean squared error (MSE), mean squared logarithmic error (MSLE) and Huber loss. **f)** The optimized iDMS network uses Glorot for weight initialization, softplus as the inner activation function, linear as the outer activation function, MAE as the loss function, and early stopping at 1E-10. **g)** Testing Phase. The optimized model was used to predict for the lipids held out in the testing dataset, benchmarked against manual derivation comparing MAE, R^2^ and P^2^, and **(h)** deployed in an LC-ESI-DMS-MS/MS method to assess “real-world” utility using a custom-synthesized standard assessing dynamic range and extent of stereoisomer cross-talk.

To solve our supervised learning problem, we employed a deep learning approach. The iDMS neural network was constructed in Python v3.12.10^18^ using the Pandas v2.3.3^19^ and NumPy v2.4.0^20^ libraries, together with the Tensorflow v2.20.0^21^, and Keras 3.13.0^22^ deep learning frameworks. Additional packages required by iDMS are listed in the requirements.txt file and full deployment instructions for Windows and MacOS is available at https://github.com/neurolipidomics/iDMS. We explored a model architecture of seven fully connected layers composed of one input layer, 5 hidden layers, and one output layer (Fig. 4c). The sequential class in Keras was used to establish the model. ReLU and softplus activation functions were evaluated for the hidden layer hyperparameters. ReLU, softplus, sigmoid, and linear activation functions were considered for the outer layer hyperparameters. Both He and Glorot uniform network initialization methods were compared in the optimization phase. We ranked combinations according to mean absolute error (MAE), mean squared error, mean squared logarithmic error, and Huber loss across all SVs for the predicted CoV µ of each stereoisomer and NI^v^ of each stereoisomer pair. All combinations of hyperparameters and initialization methods were evaluated using leave-one-lipid-pair out cross-validation (LOOV) approach (Fig. 4d,e). The Adam optimizer was used during optimization and training with a learning rate of 1E-3 for 2000 epochs and a batch size of 32. The optimized performance of the final iDMS model, employing the top-ranked hyperparameter combinations, was established using a LOOV approach on the optimization/validation dataset (Fig. 4f) before generalization ability was tested using the testing dataset (Fig. 4g). Optimized performance was benchmarked against empirical derivation comparing the *R*^2^ and MAE of replicate manual measurements to the prediction accuracy on validation data across all LOOV assessments (*P*^2^) and to the MAE of iDMS using the testing datasets, presented as parity plots. Prediction accuracy was defined as

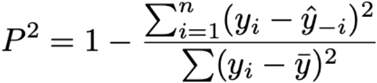

where *y_i_*= the empirical CoV, *y*^―^= the average of all CoVs in the empirical dataset, *ŷ*_-_*_i_* = the predicted value of Cov*_i_* from a model trained without seeing the validation or testing lipids, *n* =the total number of CoVs. Resolution accuracy was also benchmarked using the testing dataset to manual derivation performed as previously described^8^, comparing separability of empirically determined parameters to iDMS predicted parameters (Fig. 4g). To establish the minimal training dataset requirements, MAE was re-assessed when iDMS was trained on datasets composed of 4 to 28 monoglycosphingolipid stereoisomers (representing 2-14 stereoisomer pairs). Predictions that fell within 0.1 V of the maximal MAE and minimal (MAE/√2) boundaries of the manually determined values of each test lipid pair were considered to have met criteria for deployment. Accuracy was then tested in “real-world” application by deploying iDMS predictions in a LC-ESI-DMS-MS/MS MRM method, comparing dynamic range of custom-synthesized stereoisomers loaded both as mixtures and singly to assess iDMS-determined resolution in MRM quantification (Fig. 4h).

### iDMS Deployment Validation: LC-ESI-DMS-MS/MS MRM method

Our MRM methods were as described^8^. Briefly, LC was performed using an Agilent Infinity II High-Speed Binary LC Pump with an Agilent Infinity II system at a flow rate of 10 µl/min. The aqueous mobile phase (Solvent A) was 0.1% formic acid and 10 mM ammonium acetate. The organic mobile phase (Solvent B) was acetonitrile/isopropanol (5:2 v/v) with 0.1% formic acid and 10 mM ammonium acetate. Five µl of sample in EtOH, 2.5 µl EtOH, and 16 µl Solvent A were mixed and loaded into the autosampler at 4°C. Eight µl of this mix was injected onto an in-house packed ReproSil-Pur 120 C8 capillary column (particle size of 3 μm and pore size of 120 Å, Dr. A. Maisch, Ammerbruch, Germany) measuring 100 mm×250 µm (inner diameter). The LC gradient began at 30% Solvent B, ramped from 30% to 100% Solvent B over 5 min, was maintained at 100% Solvent B for 30 min, returned to 30% Solvent B over a 1 min period, and was re-equilibrated at 30% Solvent B for 10 min prior to the next sample or blank injection (45 min method). Each duty cycle was 0.85 s. Data acquisition was performed in positive ion MRM mode using the source and DMS parameters described in Supporting Table S2 and S3. SV and CoV were predicted using iDMS as described below.

### Software Availability

iDMS is implemented as two open-source python scripts and two Jupyter notebooks^23^. We include the two preprocessing modules as part of the DMS Data Extraction Toolkit as both python scripts and Jupyter notebooks to facilitate data processing for training set development. It is recommended that the user generate their sample datasets with CoV ramp steps of 0.2 V and deploy the optimal parameters achieved at the lowest SV in their MRM methods to extend the lifespan of their DMS (see *Optimal DMS parameters are the SV and CoV combinations that achieve quantifiable stereoisomer separation with the highest signal intensity at the lowest SV* in Results and Discussion). Module 2 outputs the resulted prediction in the csv file PredictionResult, where the predicted CoV μ for Glc and Gal isomers are listed at every SV included in the training dataset, the CoV where the isomers are calculated to intersect, predicted normalized intensity at the intersected CoV, and a classification determining whether the isomers are separated at any given SV. All scripts and notebooks are freely available at https://github.com/neurolipidomics/iDMS.

## RESULTS AND DISCUSSION

### Optimal DMS parameters are the SV and CoV combinations that achieve quantifiable stereoisomer separation with the highest signal intensity at the lowest SV

Our supervised learning problem was formulated after defining the nonlinear relationships between SV, CoV, and monoglycosphingolipid structural elements underlying FAIMS/DMS stereoisomer resolution (Fig. 5). We found that both measured and normalized signals, collected over a grid of 42 SVs and 101 CoVs in ramped steps of 0.2 V, expressed as a function of CoV followed a normal distribution at every SV for each monoglycosphingolipid stereoisomer regardless of *N*-acyl chain length or degree of unsaturation (Fig. 5a, Supporting Fig. S1). Unique Glc and Gal CoV µ, indicative of two distinct Gaussian distributions, were not observed until SVs of 2000 V and higher were applied (Fig. 5a,b). Within this range, all β-Glc stereoisomers were maximally redirected into the mass spectrometer at lower CoVs and all β-Gal stereoisomers at higher CoVs, regardless of *N*-acyl chain length or degree of saturation (Fig. 5a,b Supporting Fig. S1). Shorter *N-*acyl chains required negative and longer *N-*acyl chains required positive CoVs to compensate for increasing SV (Fig. 5a,b). The magnitude of these compensatory responses was dependent on the sugar headgroup. β-Glc stereoisomers were more responsive than β-Gal stereoisomers across all SVs when compensation required negative CoVs, meaning that decreasing compensation voltages in response to increasing SVs moved the Gaussian distribution and associated CoV µ of the β-Glc signal intensities further to the left than that of β-Gal stereoisomers (Fig. 5a,b). Conversely, β-Gal stereoisomers with longer *N*-acyl chains were more responsive than β-Glc stereoisomers when compensation required positive CoVs. Increasing compensation voltages in response to increasing SVs moved the Gaussian distribution and associated CoV µ of the β-Gal signal intensities further to the right than that of β-Glc stereoisomers (Fig. 5a,b). For β-Gal monoglycosphingolipids, dependency switched from negative to positive CoV polarity when the length of *N-*acyl chain exceeded 16 fully saturated hydrocarbons (Fig. 5a,b). The degree of unsaturation increased this inflection point. Positive CoV polarity produced maximal signal intensity when the *N*-acyl chains of β-Gal monoglycosphingolipids were composed of 22 monounsaturated hydrocarbons or more (Supporting Fig. S1). By contrast, β-Glc stereoisomers showed a greater dependency on negative CoV polarity. The inflection point did not switch until the length of *N-*acyl chain exceeded 22 fully saturated carbons or 24 monounsaturated hydrocarbons (Supporting Fig. S1).

**Figure 5.**
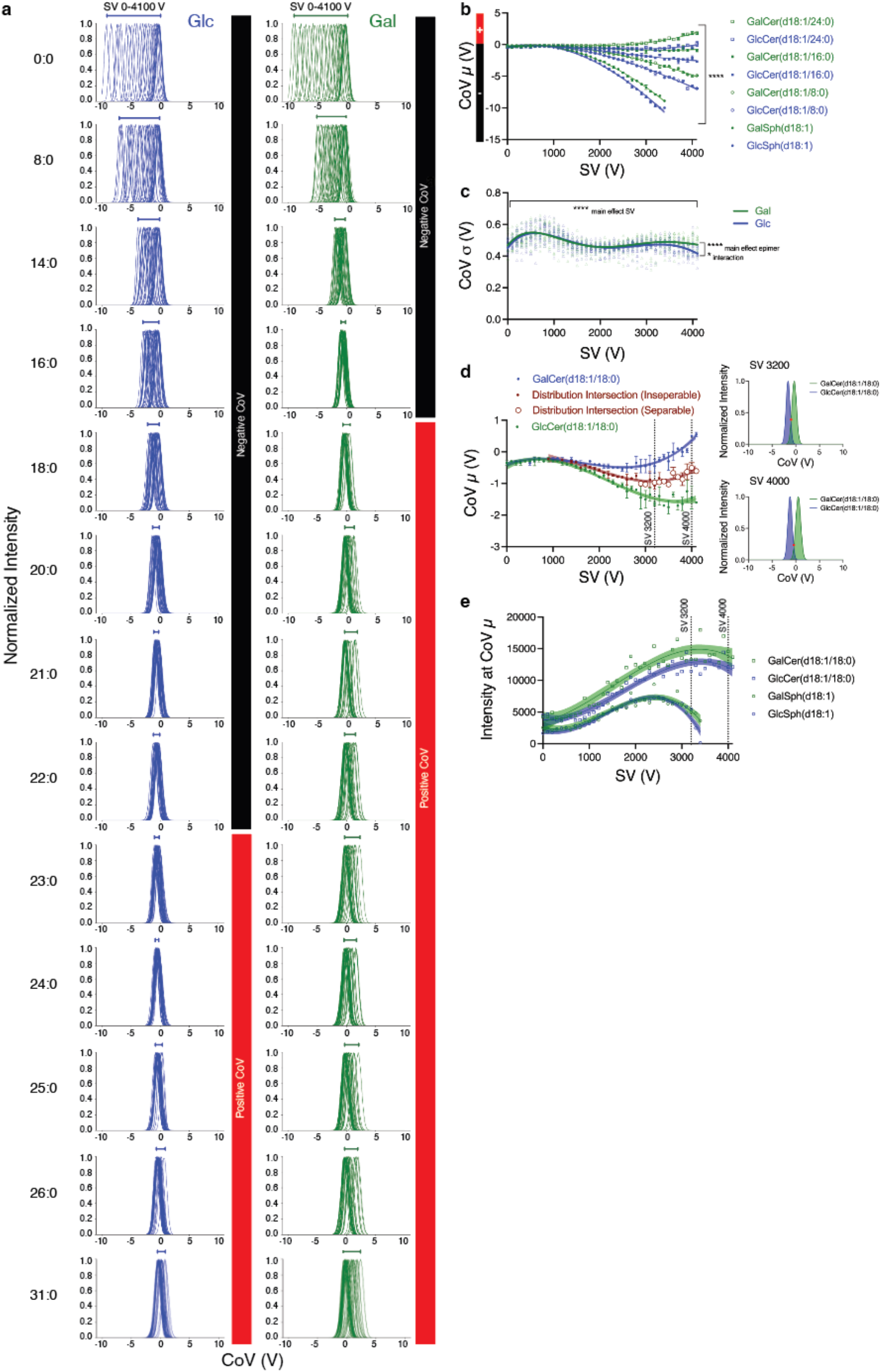
Nonlinear dependencies of CoV on SV, monoglycosphingolipid headgroup and *N-*acyl chain length. **a)** Lower CoVs are required to redirect β-Glc compared to β-Gal monosphingolipid ions to the mass spectrometer at all SVs. Shorter *N*-acyl chains require negative and longer positive CoVs regardless of headgroup. The inflection points from negative to positive CoV polarity differ by headgroup, occurring when the β-GalCer exceeds 16 and β-GlcCer *N*-acyl chain length exceeds 22 saturated hydrocarbons. Data represent normalized signal intensities expressed as a function of CoV (ionograms) at every SV (0 to 4100 V). Monounsaturated relationships are depicted in Supporting Figure S1. The inflection point in CoV polarity is indicated by the black and red bar graphs to right of the ionograms. **b)** Dispersion plots over a range of *N*-acyl chain lengths demonstrate that unique Glc and Gal CoV µ, indicative of two distinct Gaussian distributions, are not observed until SVs of at least 2000 V and higher are applied. Species with shorter *N*-acyl chain lengths require lower SVs to separate. Data represent mean CoV µ across replicate empirical experiments ± standard deviation. **c)** The σ of the normalized signal intensity distributions vary according to headgroup at higher SVs. Data represent the σ of each monosphingolipid interrogated at each SV. Line of best fit ± 95% confidence interval (CI) displays the interaction between headgroup and SV wherein the β-Glc but not β-Gal signal intensity distributions become more tightly clustered around their CoV µ at higher SVs. σ is not significantly different across all *N*-acyl chain length at each SV within headgroups but is significantly different between headgroups. Statistics: Two-way mixed effects ANOVA, main effect of SV, ****p<0.0001, SV x Lipid interaction, ****p<0.0001, main effect of Lipid, p>0.05. **d)** Resolution increases with SV. Data represent dispersions plots of replicate β-Glc and β-Gal(d18:1/18:0) experiments displaying average CoV µ ± σ and average intersection points ± σ. Solid lines indicate lines of best fit (cubic spline) ± 95% CI. Intersection points that meet the separability criteria of valley intensity < 0.5 normalized peak intensity are indicated by open circles. Insets depict the overlap of two signal intensities deemed separable (quantifiable) at 3200 and 4000 SV, also indicated by dotted lines in the dispersion plots. **e)** Higher resolution at higher SVs is associated with lower measured signal intensity. Loss is greater for isomers with shorter *N*-acyl chains. Data represent replicate measured signal intensities for β-Glc and β-Gal(d18:1/18:0) experiments. Solid lines indicate lines of best fit (cubic spline) ± 95% CI. Dotted lines map position of separation at SV 3200 and SV 4000 in **(d).**

Signal intensity distributions were dependent on sugar headgroup (Fig. 5c). The Glc σ of the Gaussian signal intensity distribution across SVs were measurably different from the Gal σ, due to a statistically significant interaction at higher SVs. Here, the β-Glc signal intensities became more tightly clustered around their CoV µ while the β-Gal distributions remained constant (Fig, 5a,c). Within epimers, intensity distributions were constant at each SV, meaning that at each SV β-Glc σ were statistically equivalent regardless of *N*-acyl chain length or degree of unsaturation as were β-Gal σ (Fig. 5c).

The relationships between resolution, SV, CoV, and ion intensity are presented as dispersion plots following the recommendations of Nolan *et al* to estimate peak resolution^24^ (Fig. 5d). Stereoisomers are considered resolved (separable) when the valley intensity (NI^v^) at the intersection point between their respective intensity distributions is less than 50% of the maximal normalized signal intensity (NI^Glcµ^, NI^Galµ^, Fig. 3b, 5d). We found, as expected, that resolution increased with higher SV (Fig. 5d); however, this improvement came at the cost of declining signal intensities (Fig. 5e). This extent of this decline was dependent on *N*-acyl chain length. Signal intensities of smaller stereoisomers with shorter *N-*acyl chains were more markedly reduced at higher SVs than species with larger *N-*acyl chains (Fig. 5e). Moreover, we found across our replicate datasets that SVs higher than 3900 V increased the risk of corona discharge resulting in high-voltage errors that ended acquisition in 30% of consecutive runs with SVs of 4000 or 4100.

This exploration enabled us to define our supervised learning problem as the prediction of CoV µ for any pair of monoglycosphingolipid stereoisomers as a function of the three most important variables impacting on resolution: 1) *N*-acyl chain length (integer feature), 2) *N*-acyl chain degree of unsaturation (integer feature) and 3) SV (integer feature).

### iDMS accurately predicts CoV µ for β-GlcCers and β-GalCers but underperforms for the monoglycosphingoid bases β-GlcSph and β-GalSph

The final iDMS neural network architecture was composed of seven fully connected layers with one input layer, five hidden layers, and one output layer (Fig. 4c). The input layer has three neurons. Layers 2-6 have ten neurons. The output layers have two neurons. The top-ranked hyperparameter combinations deployed in the optimized model are activation as the inner activation function; linear activation as the outer activation function; mean absolute error as the loss function; Glorot for weight initialization. Additional details of the model architecture are provided in Supporting Table 5.

iDMS performance was assessed using a LOOV approach, benchmarking iDMS CoV µ predictions (average MAE and P^2^) to the average MAE and co-efficient of determinations (R^2^) of empirically measured replicates across all SVs. This empirical baseline represents the maximal accuracy achievable by a predictive model before overfitting on dataset-specific technical variabilities. Table 1 demonstrates a strong overall iDMS fit for β-GlcCers and β-GalCers but not β-GlcSph and β-GalSph. Average MAEs across all species in the validation dataset fell within 0.1 V CoV of the empirically determined measures, with P^2^ approaching, but not exceeding R^2^, indicative of little to no overfitting. However, a clear bias was observed for β-GlcCers and β-GalCers. iDMS could not predict CoV µ for the monoglycosphingoid base epimers, β-GlcSph(d18:1) and β-GalSph(d18:1). P^2^ approached R^2^ yet CoV µ predictions carried a large MAE (1 and 0.64 V respectively). These voltages differences would fail to redirect ions into the mass spectrometer if deployed (Table 1). Closer examination of dispersion plots confirmed that, while the iDMS-predicted CoV µ for β-GlcCers and β-GalCers were comparable to that of empirically determined measures across all SVs and did not show any apparent bias for *N-*acyl chain length or degree of unsaturation, the predicted CoV µ for both β-Glc and β-GalSph were markedly underestimated at SVs less than 3000 V and overestimated at higher SV voltages (Supporting Fig, 2). Our attempts to further tune the iDMS hyperparameters to improve MAE for these glycosylated sphingoid bases failed. Changes resulted in a marked drop in predictive power for β-GlcCers and β-GalCers. Thus, we conclude that iDMS cannot be used to predict DMS parameters for monoglycosphingoid base epimers.

**Table 1.** Optimized iDMS model performance compared to empirical DMS parameter determination using standards.

| Lipid Identity | CoV $\mu$ (V) MAE<sup>a</sup> | | | | Coefficient of determination (CoV $\mu$)<sup>a</sup> | | | |
| --- | --- | --- | --- | --- | --- | --- | --- | --- |
| | Technical CoV $\mu$ | | iDMS Validation CoV $\mu$ | | Technical R<sup>2</sup> | | iDMS Validation P<sup>2</sup> | |
| | Glcμ | Galμ | Glcμ | Galμ | Glcμ | Galμ | Glcμ | Galμ |
| β-GlcSph(d18:1)<br>β-GalSph(d18:1) | 0.10 | 0.10 | 1.00 | 0.64 | 1.00 | 1.00 | 0.87 | 0.91 |
| β-GlcCer(d18:1/8:0)<br>β-GalCer(d18:1/8:0) | 0.11 | 0.11 | 0.19 | 0.29 | 0.99 | 0.99 | 0.98 | 0.93 |
| β-GlcCer(d18:1/14:0)<br>β-GalCer(d18:1/14:0) | 0.18 | 0.10 | 0.15 | 0.10 | 0.95 | 0.93 | 0.92 | 0.90 |
| β-GlcCer(d18:1/16:0)<br>β-GalCer(d18:1/16:0) | 0.10 | 0.13 | 0.21 | 0.23 | 0.95 | 0.65 | 0.89 | 0.10 |
| β-GlcCer(d18:1/16:1)<br>β-GalCer(d18:1/16:1) | 0.11 | 0.10 | 0.08 | 0.08 | 0.96 | 0.73 | 0.98 | 0.89 |
| β-GlcCer(d18:1/18:1)<br>β-GalCer(d18:1/18:1) | 0.15 | 0.10 | 0.11 | 0.08 | 0.85 | 0.37 | 0.92 | 0.53 |
| β-GlcCer(d18:1/20:0)<br>β-GalCer(d18:1/20:0) | 0.12 | 0.08 | 0.12 | 0.09 | 0.71 | 0.91 | 0.78 | 0.91 |
| β-GlcCer(d18:1/21:0)<br>β-GalCer(d18:1/21:0) | 0.13 | 0.12 | 0.13 | 0.09 | 0.47 | 0.90 | 0.55 | 0.93 |
| β-GlcCer(d18:1/22:0)<br>β-GalCer(d18:1/22:0) | 0.18 | 0.12 | 0.11 | 0.13 | 0.15 | 0.96 | 0.22 | 0.92 |
| β-GlcCer(d18:1/23:0)<br>β-GalCer(d18:1/23:0) | 0.13 | 0.10 | 0.11 | 0.09 | 0.56 | 0.94 | 0.58 | 0.96 |
| β-GlcCer(d18:1/24:0)<br>β-GalCer(d18:1/24:0) | 0.13 | 0.07 | 0.10 | 0.10 | 0.44 | 0.96 | 0.60 | 0.94 |
| β-GlcCer(d18:1/24:1)<br>β-GalCer(d18:1/24:1) | 0.15 | 0.09 | 0.11 | 0.17 | 0.41 | 0.92 | 0.37 | 0.62 |
| β-GlcCer(d18:1/25:0)<br>β-GalCer(d18:1/25:0) | 0.11 | 0.10 | 0.11 | 0.12 | 0.57 | 0.94 | 0.61 | 0.85 |
| β-GlcCer(d18:1/31:0)<br>β-GalCer(d18:1/31:0) | 0.11 | 0.11 | 0.16 | 0.16 | 0.79 | 0.95 | 0.32 | 0.88 |
| **Average across all lipids (Range)** | 0.13<br>(0.10-0.18) | 0.10<br>(0.07-0.13) | 0.19<br>(0.08-1.00) | 0.17<br>(0.08-0.64) | 0.70<br>(0.15-1.00) | 0.87<br>(0.37-1.00) | 0.69<br>(0.22-0.98) | 0.81<br>(0.10-0.96) |

<sup>a</sup> Technical metrics were calculated comparing replicate datasets wherein manual determination of all measures were performed using synthetic standards across all SVs tested (0-4100 V). iDMS CoV $\mu$ metrics were calculated comparing predictions to the replicate used as the training dataset.

### Generalization prediction accuracy of iDMS evaluated using the testing dataset

When trained using the training dataset of 28 lipids (14 stereoisomer pairs) across 41 SVs with a CoV ramp step of 0.2 V, the iDMS-predicted CoV µ for all six lipids fell within the 95% confidence interval of dispersion plot lines of best fit for the technical replicates, indicating an accuracy comparable to manual parameter optimization (Fig. 6a-c). Overall MAE were 0.11 V (β-GalCer(d18:1/18:0)), 0.12 V (β-GlcCer(d18:1/18:0)), 0.08 V (β-GalCer(d18:1/22:1)), 0.13 V (β-GlcCer(d18:1/22:1)), 0.09 V (β-GalCer(d18:1/26:0)), and 0.09 V (β-GlcCer(d18:1/26:0)). Model performance, examined in parity plots, confirmed precision with no obvious bias (Fig. 6d). Isomeric separability was further compared *in silico* (Fig. 6e). The onset and range of the SV and COV µ combinations meeting separability criteria predicted by iDMS matched empirically determined (Fig. 6e).

**Figure 6.**
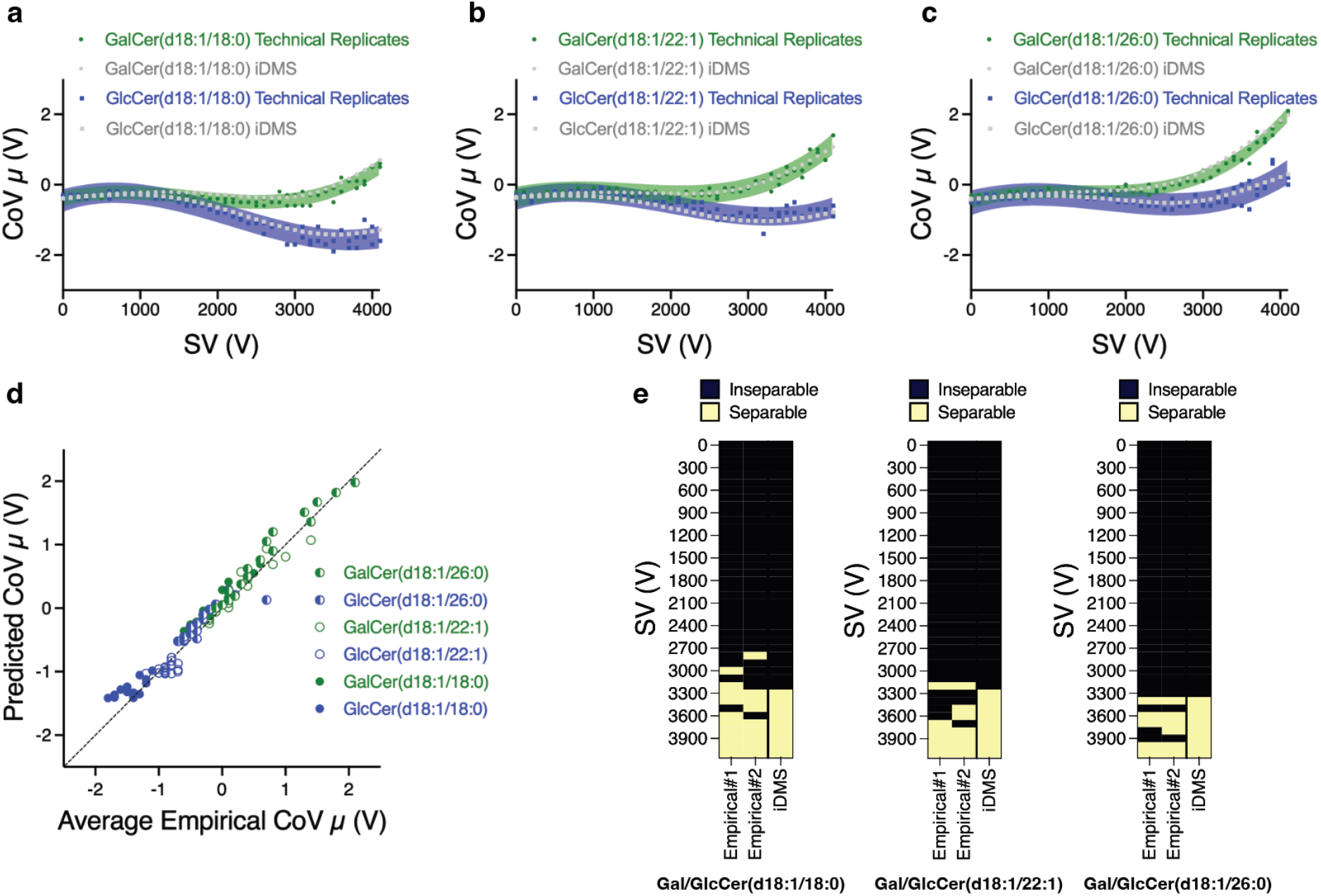
iDMS generalization prediction accuracy. Dispersion plots comparing replicate empirical determination of CoV µ to iDMS predictions of **a)** β-Glc and β-Gal(d18:1/18:0), **b)** β-Glc and β-Gal(d18:1/22:1), and **c)** β-Glc and β-Gal(d18:1/26:0). iDMS was trained using the full training dataset of 28 stereoisomers. Data display each technical replicate and lines of best fit (cubic spline) ± 95% CI. iDMS predictions fell within the empirically determined CIs. **d)** Parity plots comparing iDMS-predicted CoV µ to the average empirical CoV *μ* of the test lipids. Data represent all CoV µ values for SV 2500-4100, the SV range empirically determined in **a-c** to show two distinct β-Glc and β-Gal distributions. **e)** Classification of separability comparing iDMS to empirically determined replicate SVs and CoVs (empirical #1 and empirical #2).

### iDMS training dataset size considerations

The iDMS training set must contain, at minimum, two pairs of stereoisomers (4 lipids) composed of two different *N*-acyl chain lengths and two different degrees of unsaturation acquired across the same SV and CoV ranges to satisfy input requirements. To establish the smallest training dataset size capable of predicting CoV µ within 0.1 V of manual parameter determination, we randomly varied the training datasets ensuring these input criteria were met while predicting CoV µ using the testing dataset (Fig. 7a-c). When iDMS was trained using 6 or more pairs of monoglycosphingolipids (minimum of 12 stereoisomers), the CoV µ predictions fell within 0.1 V of the maximal MAE and minimal (MAE/√2) boundaries of the manually determined values of each test lipid pair (Fig. 7a-c). Including β-Glc/GalSph(d18:1) in the training dataset did not change generalization prediction accuracy, indicating that users can manually optimize monoglycosphingoid bases for which iDMS underperforms and include these manually determined parameters as part of their training input data. (Fig, S3).

**Figure 7.**
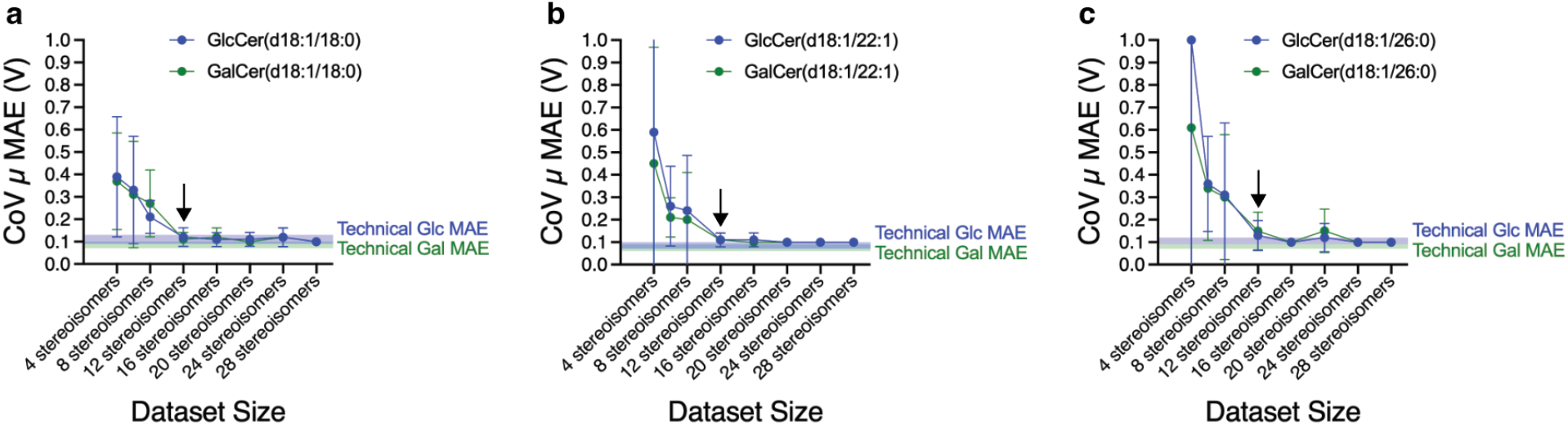
Optimal model performance is obtained using training datasets of 12 stereoisomers (6 pairs) or higher. **a)** β-Glc and β-Gal(d18:1/18:0), **b)** β-Glc and β-Gal(d18:1/22:1), and **c)** β-Glc and β-Gal(d18:1/26:0). Data represent MAE ± SD of 10 randomly generated training datasets of each size. Lines indicate the maximal MAE and minimal (MAE/√2) boundaries of the manually determined values of each test lipid pair, representing the target accuracy. Arrow indicates the smallest dataset size for which predicted CoV µ were within 0.1 V of this target.

### iDMS predicted SV and CoV µ parameters accurately resolve mixtures of monoglycosphingolipids when deployed in a LC-ESI-DMS-MS/MS MRM method

To definitively test whether iDMS returns MRM-deployable SV and CoV combinations, we predicted parameters using a randomly selected training dataset of 12 stereoisomers (6 pairs), chose iDMS-determined separable parameters at SV 3600, and deployed these parameters in a MRM LC-ESI-DMS-MS/MS method. We custom-synthesized stereoisomers for which standards were not commercially available for this test. Lipids were quantified as mixtures at concentrations ranging from 4.11 nM to 1 µM of each lipid. To determine whether these parameters effectively resoluved each epimer, 333 nM of each stereoisomer was acquired separately at the epimer-specific CoVs and SVs. The data show quantitation across the entire dynamic range of concentrations tested with R^2^ of 0.99 for both linear curves and no cross-contamination of epimeric ions at the predicted β-Glc and β-Gal CoVs (Fig. 8a,b).

**Figure 8.**
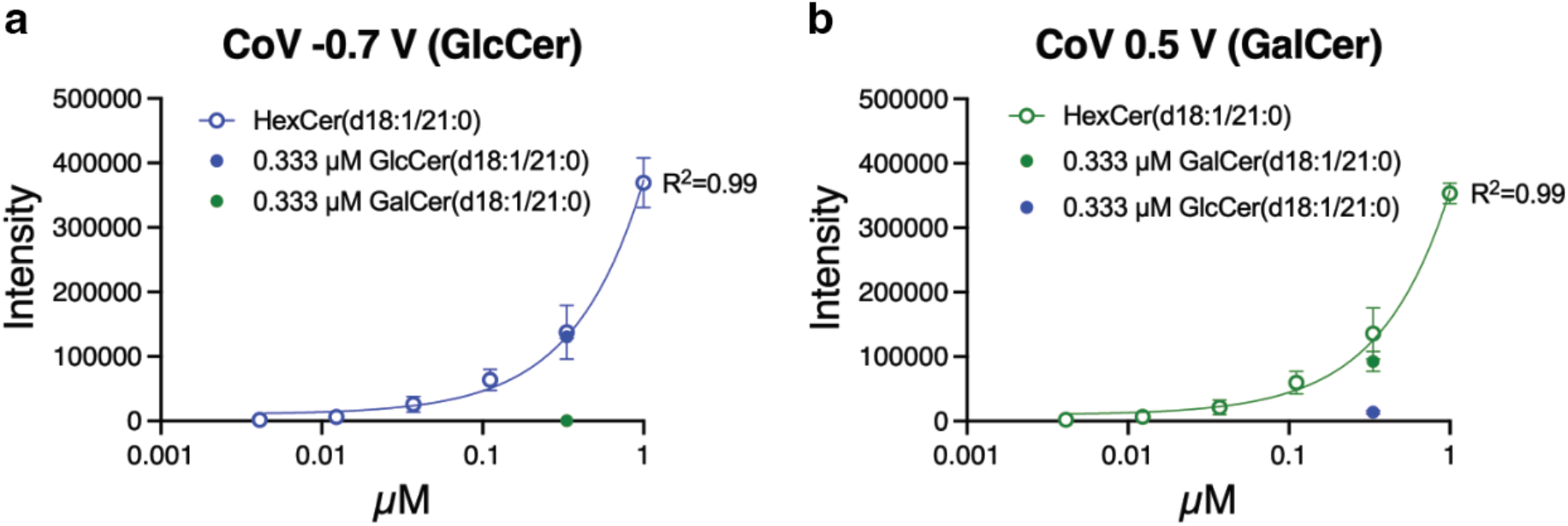
Deployment of CoV *μ* predicted by iDMS in an LC-ESI-DMS-MS/MS method. Acquisition of β-Glc and β-GalCer(d18:1/21:0) at SV3600 and **a)** CoV -0.7 V or **b)** CoV 0.5 V. Data represent mean measured signal intensity ± SD of triplicate assessments of β-Glc and β-GalCer(d18:1/21:0) loaded as a mixture (HexCer(d18:1/21:0) (open circles) or individually (filled circles). Monoglycosphingolipids were quantifiable within the tested range of 4.11 nM to 1 µM with a linear R^2^ of 0.99 at both CoVs. A CoV of -0.7 resolved of β-GlcCer(d18:1/21:0) (blue) with no detection of β-GalCer(d18:1/21:0) (green). A CoV of 0.5 resolved of β-GalCer(d18:1/21:0) (green) with little to no detection of β-GlcCer(d18:1/21:0) (blue).

## Conclusions

We developed iDMS with intent of accelerating DMS deployment in LC-ESI-DMS-MS/MS MRM and SRM methods quantifying clinically relevant monoglycosphingolipids. We show that this *in silico* supervised neural network model can learn the nonlinear relationships between SV, CoV, monoglycosphingolipid sugar headgroup, *N*-acyl chain length, and *N*-acyl degree of unsaturation required to predict CoV µ, effectively minimizing the number of manual parameter optimization experiments required to resolve comprehensive panels of monoglycosphingolipids in LC-ESI-DMS-MS/MS methods. Importantly, this machine learning approach enables, for the first time, FAIMS/DMS to be performed on stereoisomers for which a standard is not available for compound-dependent optimization. While developed and validated using monoglycosphingolipids, the iDMS neural network is compatible for use with more complex glycosphingolipid stereoisomers as long as the training dataset requirement are met. Application of iDMS promises to greatly accelerate the deployment of MRM-RPLC-ESI-DMS-MS/MS assays for the routine quantification of glycosphingolipid stereoisomers.

## ASSOCIATED CONTENT

## Data availability statement

iDMS is available as two python scripts and two modular Jupyter notebooks at https://github.com/Neurolipidomics/iDMS.

## Supporting Information

Source of monoglycosphingolipid standards used in method development; ESI source and DMS mass spec acquisition parameters; compound-dependent parameters for every transition corresponding to each pair of monoglycosphingolipid isomers used to produce training datasets; list of file and folder outputs generated by both iDMS modules; overview of iDMS neural network’s architecture; supplementary data (PDF).

Sample training dataset files and prediction list files to run iDMS and to use as templates to input user-specific training data (CSV).

## AUTHOR INFORMATION

## Corresponding Authors

**Steffany A.L. Bennett** – Neurolipidomics Laboratory and India Taylor Lipidomic Research Platform, Ottawa Institute of Systems Biology, University of Ottawa Brain and Mind Research Institute, Centre for Catalysis Research and Innovation, Department of Biochemistry, Microbiology and Immunology, Department of Cellular and Molecular, Department of Chemistry and Biomolecular Sciences, University of Ottawa, K1H 8M5, Canada. Neuroscience Program, Ottawa Hospital Research Institute, K1H 8L6, Canada. ;

**Thao Nguyen-Tran** – Neurolipidomics Laboratory and India Taylor Lipidomic Research Platform, Ottawa Institute of Systems Biology, University of Ottawa Brain and Mind Research Institute, Centre for Catalysis Research and Innovation, Department of Biochemistry, Microbiology and Immunology, Department of Cellular and Molecular, Department of Chemistry and Biomolecular Sciences, University of Ottawa, K1H 8M5, Canada. Neuroscience Program, Ottawa Hospital Research Institute, K1H 8L6, Canada. ;

## Authors

**Xun Xun Shi** – Neurolipidomics Laboratory and India Taylor Lipidomic Research Platform, Ottawa Institute of Systems Biology, Department of Biochemistry, Microbiology and Immunology, University of Ottawa, K1H 8M5, Canada

**Emily Hashimoto-Roth** – Neurolipidomics Laboratory and India Taylor Lipidomic Research Platform, Ottawa Institute of Systems Biology, Department of Biochemistry, Microbiology and Immunology, University of Ottawa, K1H 8M5, Canada; Terrence Donnelly Centre for Cellular and Biomolecular Research, Department of Molecular Genetics, University of Toronto, M5S 3K3, Canada

**Michael G Organ** – Centre for Catalysis Research and Innovation, Department of Chemistry and Biomolecular Sciences, University of Ottawa, K1N 6N5, Canada.

**Mathieu Lavallée-Adam** – Ottawa Institute of Systems Biology, Department of Biochemistry, Microbiology and Immunology, Faculty of Medicine, University of Ottawa, K1H 8M5, Canada.

**Theodore J Perkins –** Ottawa Institute of Systems Biology, Department of Biochemistry, Microbiology and Immunology, Faculty of Medicine, University of Ottawa, K1H 8M5, Canada; Regenerative Medicine Program, Ottawa Hospital Research Institute, K1H 8L6, Canada

## Author Contributions

†T.N-T., and X.X.S., contributed equally to this work. M.L-A., T.J.P,, S.A.L.B., X.X.S. and T.N-T. conceived and designed the study. T.N-T. developed the DMS methodology, wrote the Python scripts and Juypter notebooks, performed all empirical and *in silico* experiments, conducted all analyses, and wrote the manuscript under the supervision of M.G.O. and S.A.L.B.. X.X.S. established the iDMS neural network architecture, performed all hyperparameter optimizations, and wrote the initial versions of the Python scripts under the supervision of M.L-A., S.A.L.B. and T.J.P. E.H-R. reviewed, tested, and revised the final scripts and notebooks. All authors reviewed, edited, and gave approval to the final version of the manuscript.

## ACKNOWLEGEMENTS

This work was supported by a Natural Sciences and Engineering Research Council Discovery Grant (RGPIN-2019-06796), Canadian Institutes of Health Research grant (PJT 183604), and Silverstein Foundation for Parkinson’s with GBA grant to S.A.L.B and a Natural Sciences and Engineering Research Council Discovery grant to M.L-A. T.N-T. and X.X.S received Natural Sciences and Engineering Research Council CREATE Matrix Metabolomics Scholarships. T.N-T. is a Silverstein Foundation Fellow.

