## Supporting Information for "Intelligent differential ion mobility spectrometry (iDMS): A deep neural network that predicts optimal space-resolved ion mobility parameters for isomeric monoglycosphingolipids"

**Contents**: Additional details on the source of standards, instrument parameters used during collection of empirical data, iDMS outputs, neural network architecture, and supplementary data.

**Table of Contents**

Table S1. Lipid standards used in method development S3

Table S2. Electrospray source parameters and DMS settings for empirical data acquisition S4

Table S3. Compound-dependent parameters for monoglycosphingolipid isomeric pairs used to train and test iDMS S5

Table S4. iDMS outputs S6

Table S5. iDMS neural network structure S8

Figure S1 The degree of unsaturation of the *N*-acyl chain and sugar headgroup influences the negative to positive CoV polarity inflection point S9

Figure S2. iDMS accurately predicts the CoV $\mu$ redirecting maximal ion intensities of β-Glc and β-GalCers into the mass spectrometer but fails to predict β-Glc and β-GalSph(d18:1) CoV $\mu$ S10

Figure S3. Inclusion of Glc-/GalSph(d18:1) in training dataset does not affect iDMS prediction accuracy S11

**Table S1. Lipid standards used in method development**

| Lipid | Source | Catalog Number |
|---|---|---|
| β-GlcSph(d18:1) | Avanti Research | #860535 |
| β-GalSph(d18:1) | Avanti Research | #860537 |
| β-GlcCer(d18:1/8:0) | Avanti Research | #860540 |
| β-GalCer(d18:1/8:0) | Avanti Research | #860538 |
| β-GlcCer(d18:1/14:0) | Avanti Research | #860788 |
| β-GalCer(d18:1/14:0) | Avanti Research | #860785 |
| β-GlcCer(d18:1/16:0) | Avanti Research | #860539 |
| β-GalCer(d18:1/16:0) | Avanti Research | #860521 |
| β-GlcCer(d18:1/16:1) | Avanti Research | Custom Order |
| β-GalCer(d18:1/16:1) | Avanti Research | Custom Order |
| β-GlcCer(d18:1/18:0) | Avanti Research | #860547 |
| β-GalCer(d18:1/18:0) | Avanti Research | #860844 |
| β-GlcCer(d18:1/18:1) | Avanti Research | #860548 |
| β-GalCer(d18:1/18:1) | Avanti Research | #860596 |
| β-GlcCer(d18:1/20:0) | Avanti Research | #860509 |
| β-GalCer(d18:1/20:0) | Avanti Research | #860793 |
| β-GlcCer(d18:1/21:0) | Avanti Research | Custom Order |
| β-GalCer(d18:1/21:0) | Avanti Research | Custom Order |
| β-GlcCer(d18:1/22:0) | Avanti Research | #860783 |
| β-GalCer(d18:1/22:0) | Avanti Research | #860786 |
| β-GlcCer(d18:1/22:1) | Avanti Research | Custom Order |
| β-GalCer(d18:1/22:1) | Avanti Research | Custom Order |
| β-GlcCer(d18:1/23:0) | Avanti Research | #860791 |
| β-GalCer(d18:1/23:0) | Avanti Research | #860792 |
| β-GlcCer(d18:1/24:0) | Avanti Research | #860784 |
| β-GalCer(d18:1/24:0) | Avanti Research | #860854 |
| β-GlcCer(d18:1/24:1) | Avanti Research | #860549 |
| β-GalCer(d18:1/24:1) | Avanti Research | #860546 |
| β-GlcCer(d18:1/25:0) | Avanti Research | Custom Order |
| β-GalCer(d18:1/25:0) | Avanti Research | Custom Order |
| β-GlcCer(d18:1/26:0) | Avanti Research | Custom Order |
| β-GalCer(d18:1/26:0) | Avanti Research | Custom Order |
| β-GlcCer(d18:1/31:0) | Avanti Research | Custom Order |
| β-GalCer(d18:1/31:0) | Avanti Research | Custom Order |

**Table S2. Electrospray source parameters and DMS settings for empirical data acquisition**

| SOURCE PARAMETERS | |
|---|---|
| **Parameters** | **Values** |
| Curtain Gas (CUR) | 10.0 psi |
| Collision Gas (CAD) | 10.0 psi |
| Ionization Voltage (IS) | 5500.0 V |
| ESI Source Temperature (TEM) | 250.0°C |
| Ion Source Gas1 (GS1) | 25.0 psi |
| Ion Source Gas2 (GS2) | 20.0 psi |
| **DMS PARAMETERS** | |
| **Parameters** | **Values** |
| DMS Temperature (DT) | Low, 150°C |
| DMS Resolution Enhancement (DR) | Medium, 20 msec |
| Modifier (MD) | 2-Propanol |
| Modifier Composition (MDC) | Low, 1.5% |
| Modifier Density (MDD) | 0.7860 g/mL |
| Modifier Weight (MDW) | 60.10 g/mol |
| DMS Offset (DMO) | -3.0 V |
| Dwell Time | 50.0 msec |

**Table S3. Compound-dependent parameters for monoglycosphingolipid isomeric pairs used to train iDMS**

| Lipid | Q1 | Q3 | DP (V) | EP (V) | CE (V) | CXP (V) |
|---|---|---|---|---|---|---|
| β-GlcSph(d18:1) and β-GalSph(d18:1) | 462.5 | 264.3 | 186 | 10 | 35 | 20 |
| β-GlcCer(d18:1/8:0) and β-GalCer(d18:1/8:0) | 588.8 | 264.3 | 271 | 10 | 45 | 38 |
| β-GlcCer(d18:1/14:0) and β-GalCer(d18:1/14:0) | 672.5 | 264.3 | 86 | 10 | 51 | 38 |
| β-GlcCer(d18:1/16:0) and β-GalCer(d18:1/16:0) | 700.7 | 264.3 | 66 | 10 | 47 | 18 |
| β-GlcCer(d18:1/16:1) and β-GalCer(d18:1/16:1) | 698.5 | 264.3 | 111 | 10 | 45 | 18 |
| β-GlcCer(d18:1/18:0) and β-GalCer(d18:1/18:0) | 728.7 | 264.3 | 136 | 10 | 45 | 32 |
| β-GlcCer(d18:1/18:1) and β-GalCer(d18:1/18:1) | 726.6 | 264.3 | 256 | 10 | 45 | 32 |
| β-GlcCer(d18:1/20:0) and β-GalCer(d18:1/20:0) | 756.6 | 264.3 | 256 | 10 | 53 | 20 |
| β-GlcCer(d18:1/21:0) and β-GalCer(d18:1/21:0) | 770.7 | 264.3 | 256 | 10 | 49 | 20 |
| β-GlcCer(d18:1/22:0) and β-GalCer(d18:1/22:0) | 784.7 | 264.3 | 251 | 10 | 59 | 20 |
| β-GlcCer(d18:1/22:1) and β-GalCer(d18:1/22:1) | 782.9 | 264.3 | 256 | 10 | 47 | 22 |
| β-GlcCer(d18:1/23:0) and β-GalCer(d18:1/23:0) | 798.7 | 264.3 | 256 | 10 | 51 | 22 |
| β-GlcCer(d18:1/24:0) and β-GalCer(d18:1/24:0) | 812.7 | 264.3 | 256 | 10 | 53 | 6 |
| β-GlcCer(d18:1/24:1) and β-GalCer(d18:1/24:1) | 810.8 | 264.3 | 246 | 10 | 51 | 22 |
| β-GlcCer(d18:1/25:0) and β-GalCer(d18:1/25:0) | 826.9 | 264.3 | 241 | 10 | 55 | 20 |
| β-GlcCer(d18:1/26:0) and β-GalCer(d18:1/26:0) | 840.8 | 264.3 | 246 | 10 | 53 | 22 |
| β-GlcCer(d18:1/31:0) and β-GalCer(d18:1/31:0) | 911.0 | 264.3 | 256 | 10 | 55 | 20 |

**Table S4: iDMS outputs**

| Module | File or Folder Name | File Format | Contents |
|---|---|---|---|
| | Lipid_data | .csv | Identity of isomers from training dataset (eg. Glc and Gal), backbone (eg. d18:1), and the list of lipids present in training dataset. |
| **Module1_Parametric Modelling** | CombinedAnalysis | .csv | Lipid species, SV, sphingoid backbone information, *N*-acyl chain length, degree of unsaturation, Gaussian-derived μ and Gaussian-derived σ at each SV for each isomer, extracted signal intensity at Gaussian-derived μ for each isomer. |
| | extractGlcMaxCoV<br>extractGalMaxCoV | .csv | Max signal intensity and corresponding CoV at each SV, *N*-acyl chain length, degree of unsaturation, replicate number, isomer identification, sphingoid backbone information, lipid species |
| | Intersected_gauss | .csv | Index, lipid species, lipid identifier, SV, *N*-acyl chain length, degree of unsaturation, Gaussian-derived μ, Gaussian-derived σ at each SV, extracted signal intensity at Gaussian-derived μ, intersection points, normalized intensity at intersection point, classification of separability at μ CoV for each SV. |
| | meanAllsigma | .csv | Average σ across all CoVs at each SV. |
| | norm_Glc<br>norm_Gal | .csv | Signal intensities, intensities normalized to max signal intensity at each CoV for every SV, *N*-acyl chain length, degree of unsaturation, sphingoid backbone information, lipid species |
| | norm_GlcParameters<br>norm_GalParameters | .csv | Signal intensity and normalized intensity CoV of max signal for each SV, *N*-acyl chain length, degree of unsaturation, isomer identification, sphingoid backbone information, lipid species, empirical μ, Gaussian-derived μ, Gaussian-derived σ, $R^2$ value of Gaussian fitting. |
| | python_version | .txt | Output of information related to the python version used to run Module 1. |
| | Plots -> ExtractIonograms (folder) | .png | Extracted ionograms |
| | Plots -> NormIonograms (folder) | .png | Normalized ionograms |

| Module | File or Folder Name | File Format | Contents |
| --- | --- | --- | --- |
| | Plots -> Glc_GaussianPlots (folder)<br>Plots -> Gal_GaussianPlots (folder) | .png | Gaussian ionograms |
| | Plots -> GaussianIntersectPlots (folder) | .png | Combined Glc and Gal Gaussian ionograms indicating intersection points. |
| **Module2 – iDMS** | iDMS_params | .csv | List of configurations associated to training iDMS. These include the verbosity, number of epochs, and total number of training steps executed per epoch used within each execution of module 2. |
| | iDMS_training_history | .csv | List of loss values and accuracy metric recorded at the end of each epoch executed during module 2. |
| | PredictionResult | .csv | Index, lipid species, SV, *N*-acyl chain length, degree of unsaturation, predicted μ for each of Gal or Glc isomer, Gaussian-derived σ at each SV, intersection points, valley CoV, normalized intensity at intersection point, classification of separability at μ CoV for each SV. |
| | TrainingSet | .csv | Data table of Glc and Gal sphingolipid isomers used as training dataset, obtained from Module1 outputs. Containing lipid species, SV, sphingoid backbone information, *N*-acyl chain length, degree of unsaturation, Gaussian-derived μ and Gaussian-derived σ at each SV for each isomer, extracted signal intensity at Gaussian-derived μ for each isomer. |
| | Python_version | .txt | Output of information related to the python version used to run Module 1. |
| | PredictedSetPlots (folder) | .png | Ionograms of Glc and Gal sphingolipid isomers based on predicted CoV μ at each SV, with indication of intersection point (if any). |

**Table S5: iDMS Neural Network Structure**

| Layer (type) | Output Shape | Param # |
|---|---|---|
| dense (Dense) | (None, 10) | 40 |
| dense_1 (Dense) | (None, 10) | 110 |
| dense_2 (Dense) | (None, 10) | 110 |
| dense_3 (Dense) | (None, 10) | 110 |
| dense_4 (Dense) | (None, 10) | 110 |
| dense_5 (Dense) | (None, 2) | 22 |

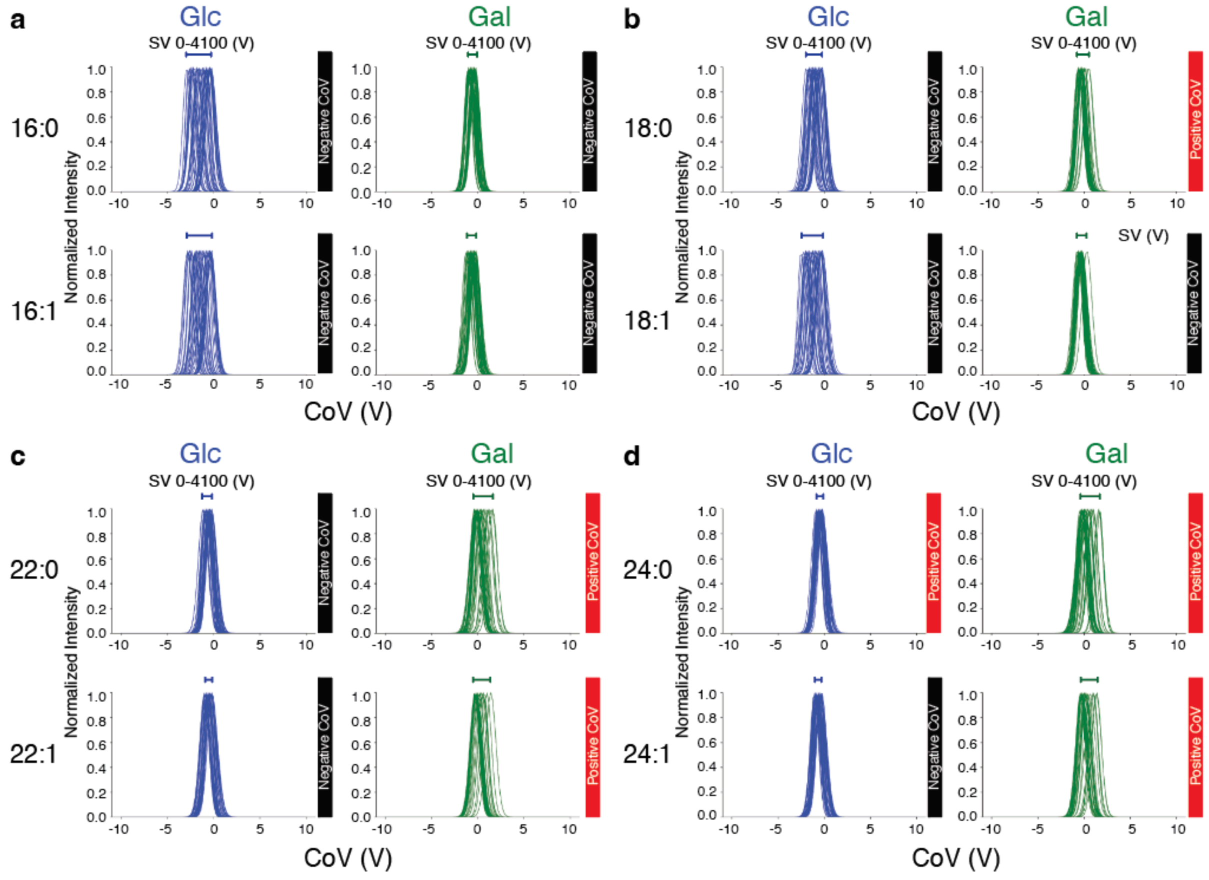


**Figure S1. The degree of unsaturation of the *N*-acyl chain and sugar headgroup influences the negative to positive CoV polarity inflection point. a)** β-Glc and β-Gal(d18:1/16:0) and (d18:1/16:1). **b)** β-Glc and β-Gal(d18:1/18:0) and (d18:1/18:1). **c)** β-Glc and β-Gal(d18:1/22:0) and (d18:1/22:1). **d)** β-Glc and β-Gal(d18:1/24:0) and (d18:1/24:1). Data represent normalized signal intensities expressed as a function of CoV at every SV (0 to 4100 V). CoV polarity is indicated by black and red bar graphs to right of the ionograms. Note that monounsaturation of the N-acyl chain increases the hydrocarbon chain length of this inflection point from 18 to 22 hydrocarbons for β-Gal stereoisomers and from 23 to greater than 24 hydrocarbons for β-Glc stereoisomers.

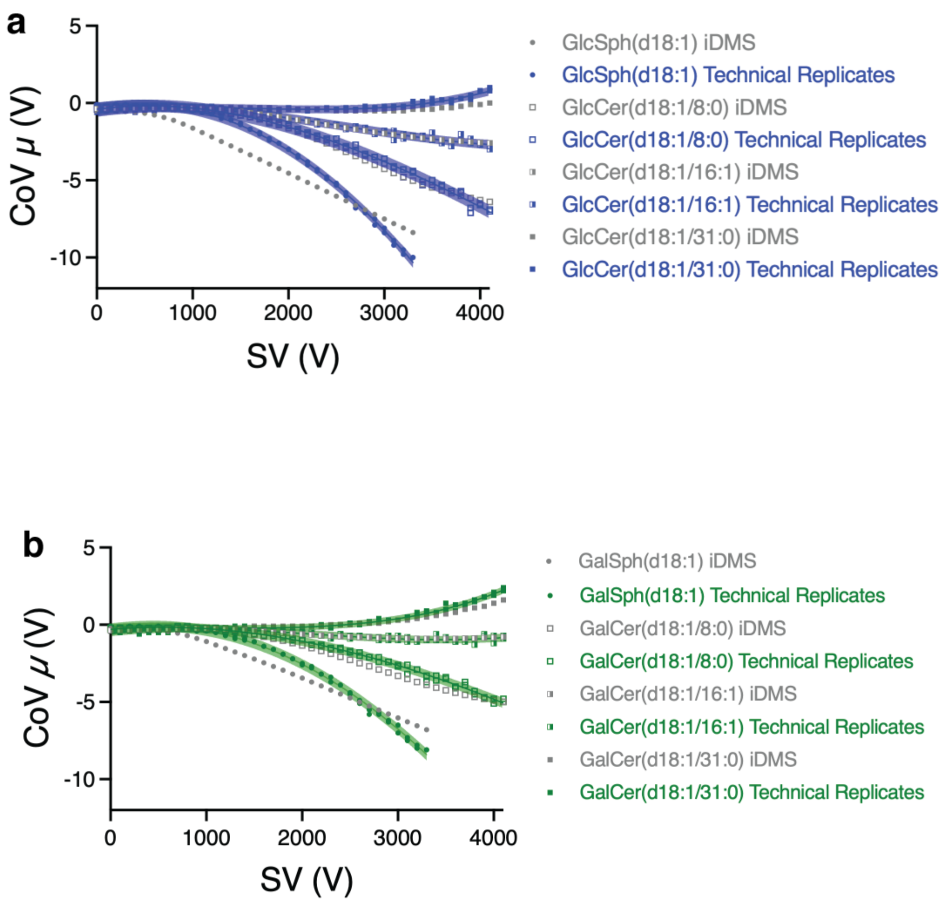


**Figure S2. iDMS accurately predicts the CoV $\mu$ redirecting maximal ion intensities of β-Glc and β-GalCers into the mass spectrometer but fails to accurately predict β-Glc and β-GalSph(d18:1) CoV $\mu$. a)** β-Glc stereoisomers. **b)** β-Glc stereoisomers. Data are presented as dispersion plots graphing empirically derived replicate CoV μ and iDMS predictions following LOOV. Solid lines represent the lines of best fit (cubic spline) $\pm$ 95% CI of the technical replicates.

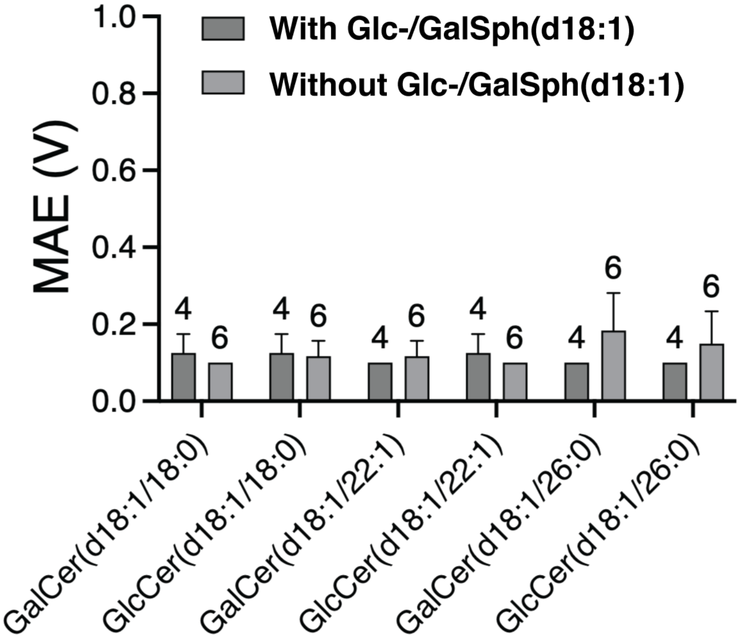


**Figure S3. Inclusion of Glc-/GalSph(d18:1) in training dataset does not affect iDMS prediction accuracy**. iDMS cannot accurately predict CoV μ when trained on a dataset composed exclusively of monoglycoceramides (see Table 1). Including Glc-/GalSph(d18:1) in the minimal training dataset of 12 lipids (6 pairs) does not alter prediction accuracy of monoglycoceramides. Data represent average MAE ± SD for each test lipid across the entire SV range (0-4100) with or without Glc-/GalSph(d18:1).
